# Age-dependent expression and antiviral activity of interferon epsilon in respiratory epithelium

**DOI:** 10.1101/2025.04.08.647778

**Authors:** Mary McCabe, Helen E. Groves, Erin Getty, Emma Campbell, Connor G.G. Bamford, Guillermo Lopez Campos, Michael D. Shields, Ultan F. Power

## Abstract

Respiratory syncytial virus (RSV) disease burden is greatest between six weeks and 6 months of life, with young age the most common risk factor among hospitalised children. A robust innate immune response in the airway epithelium is crucial for mitigating RSV-associated disease, but early-life immune responses to infection remain largely unexplored. RNA-seq analysis of RSV-infected primary nasal epithelial cell cultures from healthy infants at birth and one year revealed diminished expression of interferon epsilon (*IFNE*), a poorly characterised type I IFN, in newborns versus one-year. We hypothesised, therefore, that *IFNE* plays an important role during infant RSV infection. We found that *IFNE* is endogenously expressed in airway epithelial cell lines. Recombinant human IFNε (rhIFNε) induced an antiviral state against an RSV clinical isolate and related Sendai virus, but not SARS-CoV-2 under the conditions tested. The antiviral potency of rhIFNε was diminished relative to rhIFN beta (rhIFNβ1) or rhIFN lambda-1 (rhIFNλ1), as evidenced by IC_50_ data. Importantly, rhIFNε induced similar ISGs as rhIFNβ1 but demonstrated a transient temporal expression profile that differed from both rhIFNβ1 and rhIFNλ1. These results suggest that lower IFNε expression at birth may contribute to increased susceptibility to severe RSV-associated disease, offering insights into potential therapeutic interventions.

## Introduction

Respiratory syncytial virus (RSV), a single-stranded negative-sense RNA virus of the *Pneumoviridae* family and *Orthopneumovirus* genus, is the commonest cause of acute lower respiratory tract infection (ALRTI) in infants globally [1]. Annually, ∼33 million RSV LRTIs occur in children <5 years old, resulting in ∼101,400 infant deaths [1]. Despite recent pharmacological advancements [2–6], treatment for severe RSV-associated disease remains palliative, with no RSV-specific antivirals or direct vaccination available for infants <6 months, the age group in which the disease burden is greatest [1, 7]. While epidemiological data has identified risk factors [8–14], most infants hospitalised share only the risk factor of age, suggesting host intrinsic features predispose them to severe disease. Because of the current incomplete understanding of RSV-host interactions in early life, we cannot predict which infants will experience severe disease and, thereby, prioritise prophylactic treatment. As RSV initially targets the ciliated cells of the nasopharynx [15–17], efficient innate immune responses play a vital role in determining RSV infection outcomes. Yet, whether and how these responses develop over time, and why infants <6 months are more frequently susceptible to severe disease, is unknown.

Interferons (IFN) are a crucial innate barrier against viral infection and dissemination [18, 19]. In the respiratory epithelium, type I (IFNα and -β) and type III (IFNλ1-λ4) IFNs play significant roles. Recently, Taveras *et al,* determined that all IFN types (I, II and III) from nasal samples were significantly increased in children (>6 years) versus infants (<6 months) and higher in milder outpatient RSV infections compared to severe inpatient infections [20]. Furthermore, we observed a threefold increase in mean IFNλ1 protein secretion at one year versus birth within paired birth and one-year primary nasopharyngeal cell samples (unpublished data, Groves HE, McCabe M, Coey J, Broadbent L, Guo-Parke H, Lopez Campos G, et al). These results suggest that strong IFN responses are a critical component of determining disease severity. As such, more robust IFN responses to RSV infection with increasing chronological age may contribute to protection against severe disease in older infants and adults.

IFN epsilon (*IFNE*/IFNε) is a highly conserved type I IFN recognised for its multifaceted antimicrobial properties in the female reproductive tract (FRT) and its expression at other mucosal interfaces, including the lungs [21–24]. In contrast to other type I IFN subtypes, IFNε is 100-1000-fold less potent, and is expressed at high levels basally [25–27]. Interestingly, Thwaites *et al* described a trend for increased expression of type I IFNs (*IFNA1*, *IFNB1* and *IFNE*) in infants with moderate RSV ALRTI relative to severe cases [28] and *IFNE* was upregulated in children and young adults with SARS-CoV-2 infection [29]. Furthermore, *IFNE* was observed to be differentially expressed in children with long-coronavirus disease (COVID) symptoms [30]. Therefore, despite its conserved mucosal antiviral activity, understanding of the local expression and function of IFNε in airway epithelium against respiratory viruses, and its possible contributions to viral disease severity, remain limited.

In this study, we identified for the first time *IFNE* as differentially expressed with chronological age and confirmed its endogenous expression in immortalised airway epithelial cells. We assessed its antiviral activity against a clinical isolate of RSV and a related RNA virus, Sendai virus, and SARS-CoV-2. We determined the IC_50_ concentration of rhIFNε against RSV compared to other relevant mucosal IFNs and characterised the induction of downstream interferon-stimulated genes (ISGs). These results suggest an underappreciated role of IFNε in the induction of protective antiviral responses in the respiratory epithelium.

## Materials and Methods

### Cell Culture

BEAS-2B cells (kindly provided by Cliff Taggart, Queen’s University Belfast) were maintained in 1 g/L (low) glucose DMEM with 10 units/mL Pen/Strep and 5% FBS. HEp-2 cells (kindly provided by Prof. Ralph Tripp, University of Georgia) were maintained in 4.5 g/L (high) glucose DMEM with 10 units/mL Pen/Strep and 5% FBS. A549, Vero E6 and Vero’s expressing ACE2 and TMPRSS2 (VAT) cells were maintained in 4.5 g/L (high) glucose DMEM with 10 units/mL Pen/Strep and 10% FBS. A549 expressing human ACE2 and TMPRSS2 (AAT) cells were maintained in 4.5 g/L (high) glucose DMEM with 10 units/mL Pen/Strep, 10% FBS and maintained under antibiotic selection with Hygromycin B (25 ug/mL) and Geneticin (25 ug/mL).

Nasal brushings were performed on healthy newborn term infants, (37–42 weeks gestation) and preterm infants (28–34 weeks gestation) at birth and repeated in the same infants at one-year-old, as previously described [31]. Harvested paediatric nasal cells were cultured to differentiation using established methods [16][32] and infected with Multiplicity of Infection (MOI) 3 in duplicate with a clinical isolate of RSV, designated BT2a, or mock-infected as previously described [33].

### RNA-Seq

Total RNA was extracted from WD-PNEC cultures at 96 hpi (High pure RNA isolation kit, Roche diagnostics, Indianapolis, USA) and RNA quality was determined (Agilent RNA 6000 Nano Kit) using the 2100 Bioanalyzer Instrument (Agilent Technologies). Total RNA sequencing was conducted (unpublished data, Groves HE, McCabe M, Coey J, Broadbent L, Guo-Parke H, Lopez Campos G, et al).

### Virus Stocks and Titrations

RSV BT2a (clinical isolate) was isolated and characterised as previously described [33]. RSV A2/mkate2 (a recombinant RSV reporter virus expressing Katushka, a far-red fluorescent protein) was rescued from an infectious clone and helper plasmids kindly provided by Dr Martin Moore (Emory University, Atlanta, USA). Sendai virus/eGFP (rSeV/eGFP) was generated by reverse genetics [34]. SARS-CoV-2 Delta Variant was isolated from nasal swabs [35]. EMCV-Rueckert was obtained as described previously [36]. Viral infections were performed at the MOI stated in each figure legend by adding diluted virus suspensions to the apical compartment of WD-PNECs or directly onto cell lines in serum-free medium. HEp-2 cells were used to titrate RSV BT2a, and Vero E6 cells were used to titrate EMCV by tissue culture infectious dose 50 (TCID_50_) assays. SARS-CoV-2 plaque assays were performed on VAT cells [35].

### RNA extraction and RT-qPCR

RNA was extracted according to the manufacturer’s instructions (High pure RNA isolation kit, Roche) and RNA quality was determined (Nanodrop One, Thermo Scientific®). cDNA synthesis was performed in a 96-well thermocycler (Bio-Rad) using a High-Capacity cDNA Reverse Transcription Kit (Applied Biosystems), according to the manufacturer’s instructions. For the target genes, *IFNE* (assay ID Hs00703565_s1*), IFNL1* (assay ID Hs00601677_g1), *IFNB1* (assay ID Hs01077958_s1)*, MX1* (assay ID Hs00895608_m1)*, ISG15* (assay ID Hs01921425_s1)*, IFIT1* (assay ID Hs03027069_s1)*, RSDA2* (assay ID Hs00369813_m1)*, IRF1* (assay ID Hs00971965_m1), *CXCL10* (assay ID Hs00171042_m1), YWHAZ (assay ID Hs01122445_g1), TBP (assay ID Hs00427620_m1) and IPO8 (assay ID Hs00914057_m1) Thermo Fischer Scientific TaqMan ready-to-use primer/probes were purchased and employed with the LightCycler 480 Probes Master Mix (Roche). Relative quantification analysis of gene expression was performed using LightCycler 96 software (Roche). The Relative Quantification Cycle (Cq) of the sample was determined by comparing the target gene to 3 reference housekeeping genes: *YWHAZ*, *TBP*, and *IPO8*.

### Recombinant human protein

Cell lines were pretreated with recombinant human interferon epsilon (rhIFNε) (R&D systems, Minneapolis, MN, USA), rhIFN beta (rhIFNβ1) (Peprotech®) and rhIFN lambda-1 (rhIFNλ1) (Peprotech®) in low-serum DMEM for intervals stated within each figure legend. Cells were then infected in serum-free DMEM. Following infection, cells were incubated for 2 h at 37 °C. After 2 h, the viral inoculum was removed and washed three-times with serum-free DMEM to remove unattached virus, low-serum DMEM replaced and incubated at 37°C.

### Cell Imaging and Analysis

The mean % fluorescence ratio of RSV A2/mkate2 and rSeV/eGFP infected cells was measured using the brightfield and red/green fluorescence channels of the Celigo Imaging Cytometer (Nexcelom). All data were presented as % fluorescence ratio ((brightfield confluence/red or green confluence) *100).

### Generation of BEAS-2B IFNLR1^-/-^ cells

BEAS-2B cells overexpressing Cas9 were generated via lentiviral transduction of a third generation lentiviral Cas9 expression plasmid (Addgene #52962), blasticidin selection (10 ug/mL) and single-cell clonal isolation. Western blotting was used to confirm the stable expression of Cas9 (Sigma-Aldrich, F3165). To generate *IFNLR1* BEAS-2B clonal cell line knockouts, a single-guide RNA (sgRNA) (priCRISPR.IFNLR1.1(+) CACCGGTATTCGGACTCCACCCAG) was designed. For each sgRNA, a forward and reverse oligonucleotide sequence was synthesised (Eurofins), annealed and inserted into the lentiGuide-Puro vector via BsmB1-v2 (NEB) restriction digestion and T4 DNA ligase (NEB) mediated ligation. Vector-sgRNA DNA was then transformed into DH10B competent cells (Thermo Fischer Scientific) and spread onto Ampicillin selection plates (50 mg/mL). Single colonies were picked and miniprepped using PureLink™ HiPure Plasmid Miniprep kit according to manufacturer’s instructions (Invitrogen™). BEAS-2B *IFNLR1* knockout cell lines were generated via lentiviral transduction of a third generation lentiGuide-Puro plasmid (Addgene #84752) with inserted sgRNA sequence, puromycin (Gibco) selection and single-cell clonal isolation.

### Statistical Analysis

Statistical analyses included an unpaired two-tailed Student’s t-test and one-way analysis of variance (ANOVA) to assess the significance of test conditions, as stated in the figure legends. For the time course profiles we calculated the Area under Curve (AUC) and then used this summary measure for comparisons using one-way ANOVA. Statistical analysis was set at * p<0.05, ** p<0.01, ***p<0.001, ****p<0.0001. Data presented as means ± standard error of the means (SEM). Data were analysed using GraphPad® Prism 10 (GraphPad Software, Inc, La Jolla, CA).

### Data Availability

RNA-seq data has been deposited under moratorium pending publication and will be made available to reviewers upon request. The GEO accession number for the RNA-seq data is: GSE231666.

## Results

### Endogenous expression of *IFNE* increases during the first year of life

In a recent study, we evaluated the impact of gestational and chronological age on RSV-induced airway epithelium innate immune responses using RNA-Seq (unpublished data, Groves HE, McCabe M, Coey J, Broadbent L, Guo-Parke H, Lopez Campos G, et al) (**Figure 1A**). RNA-seq analysis of RSV-infected WD-PNECs derived from infants sampled at birth and one-year-old revealed 63 differentially expressed genes (DEGs) (42 upregulated, 21 downregulated) with Log2Fold >|2| when comparing RSV-induced responses from 1-year-versus newborn-derived cultures (**Figure 1B**). Included in the 42 upregulated DEGs, *IFNE,* a poorly characterised type I IFN, was significantly increased following RSV infection versus uninfected controls (**Figure 1C**). Furthermore, *IFNE* expression was higher in 1 year-versus newborn-derived WD-PNECs, both at baseline and following RSV infection (**Figure 1C**). These results suggested that *IFNE* may play an unappreciated antiviral role in respiratory airway epithelium and contribute to more robust antiviral activities at 1 year compared to newborns.

**Figure 1.**
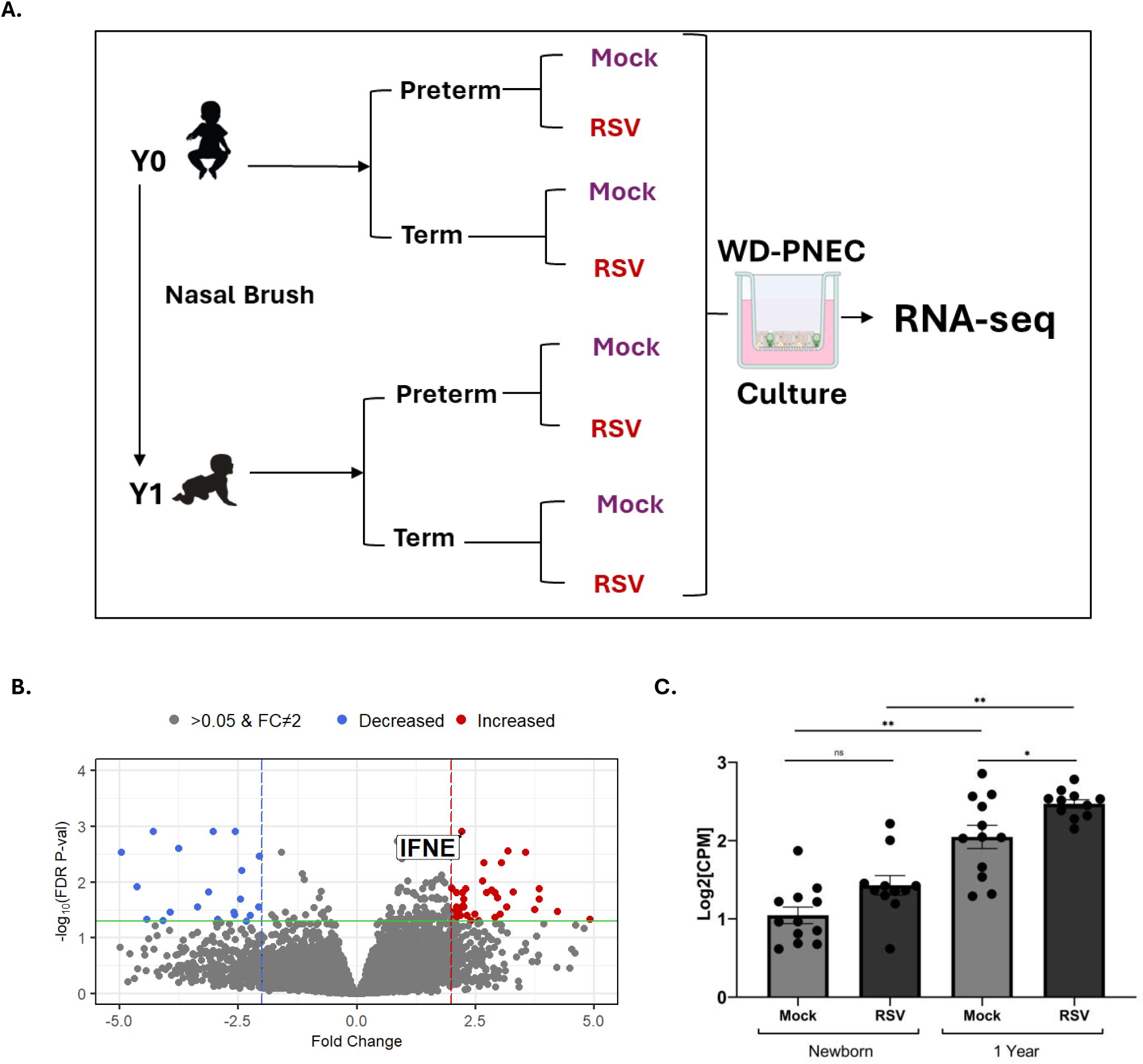
Increase in *IFNE* expression during the first year of life. (**A**) Summary of RNA-Seq experimental set-up. WD-PNEC cultures (n=12 paediatric donors) were infected with clinical isolate RSV BT2a (MOI=3) or mock-infected. RNA was extracted at 96 hpi and total RNA sequencing was conducted. (**B**) Volcano plot displaying differentially expressed genes associated with RSV-infected cultures from 1-year-old and newborn donors, with *IFNE* expression highlighted. X-axis depicts the log2 differences in gene expression for each comparison of interest, with positive values (in red) displaying significantly increased gene expression (log2 FC ≥ 2) and negative values (in blue) displaying significantly decreased gene expression (log2 FC ≤ −2). Y axis depicts adjusted p values (-log10 scale) for each gene (higher p values representing greater statistical significance) after adjusting for batch by the year RNA-sequencing was completed. Green horizontal line indicates 0.05 p-value significance cut-off (**C**) *IFNE* Log2 Counts per Million (CPM). Statistical analyses were corrected for multiple testing by applying the Benjamini-Hochberg (BH) p value correction (* p<0.05, ** p<0.01).

### Endogenous expression of *IFNE* in immortalised airway epithelial cells

Next, we assessed the expression of *IFNE* and two other mucosal IFNs within airway epithelial cells, including a type I (*IFNB1)* and a type III IFN (*IFNL1)*. *IFNB1* was chosen as a comparator as it was the sole type I IFN differentially expressed with infection within the RNA-seq data (unpublished data, Groves HE, McCabe M, Coey J, Broadbent L, Guo-Parke H, Lopez Campos G, et al). *IFNλ1* was chosen as a representative type III IFN as our group showed increased IFNλ1 release from primary cultures derived from infant respiratory tract samples following RSV infection [37]. Within common airway epithelial cell lines used to model RSV infection, *IFNE* was constitutively expressed at low levels, with the highest expression in adenocarcinoma human alveolar basal cells (A549) and A549 cells over-expressing human ACE2 and TMPRSS2 (AAT) (**Figure 2A**). Low levels of expression were evident in immortalised human bronchial epithelial cells (BEAS-2B) and human laryngeal carcinoma cells (HEp-2) (**Figure 2A**). This contrasted with *IFNB1* and *IFNL1*, which exhibited negligible expression under unstimulated sterile cell-culture conditions (**Figure 2B-C**). To further characterise *IFNE* expression, we infected A549 (**Figure 2D-F**) and BEAS-2B (**Figure 2G-I**) cells with a recombinant RSV reporter virus expressing a far-red fluorescent protein (RSV-A2/mKate2) or mock-infected them. At 48 hpi *IFNE*, *IFNB1* and *IFNL1* mRNA expression was higher within A549 cells (**Figure 2E-F**) compared to BEAS-2B cells (**Figure 2H-I**). *IFNE* expression was detected irrespective of infection in both cell lines (**Figure D, G**), with this unique endogenous pattern emphasized when comparing its expression to that of *IFNB1* or *IFNL1*.

**Figure 2.**
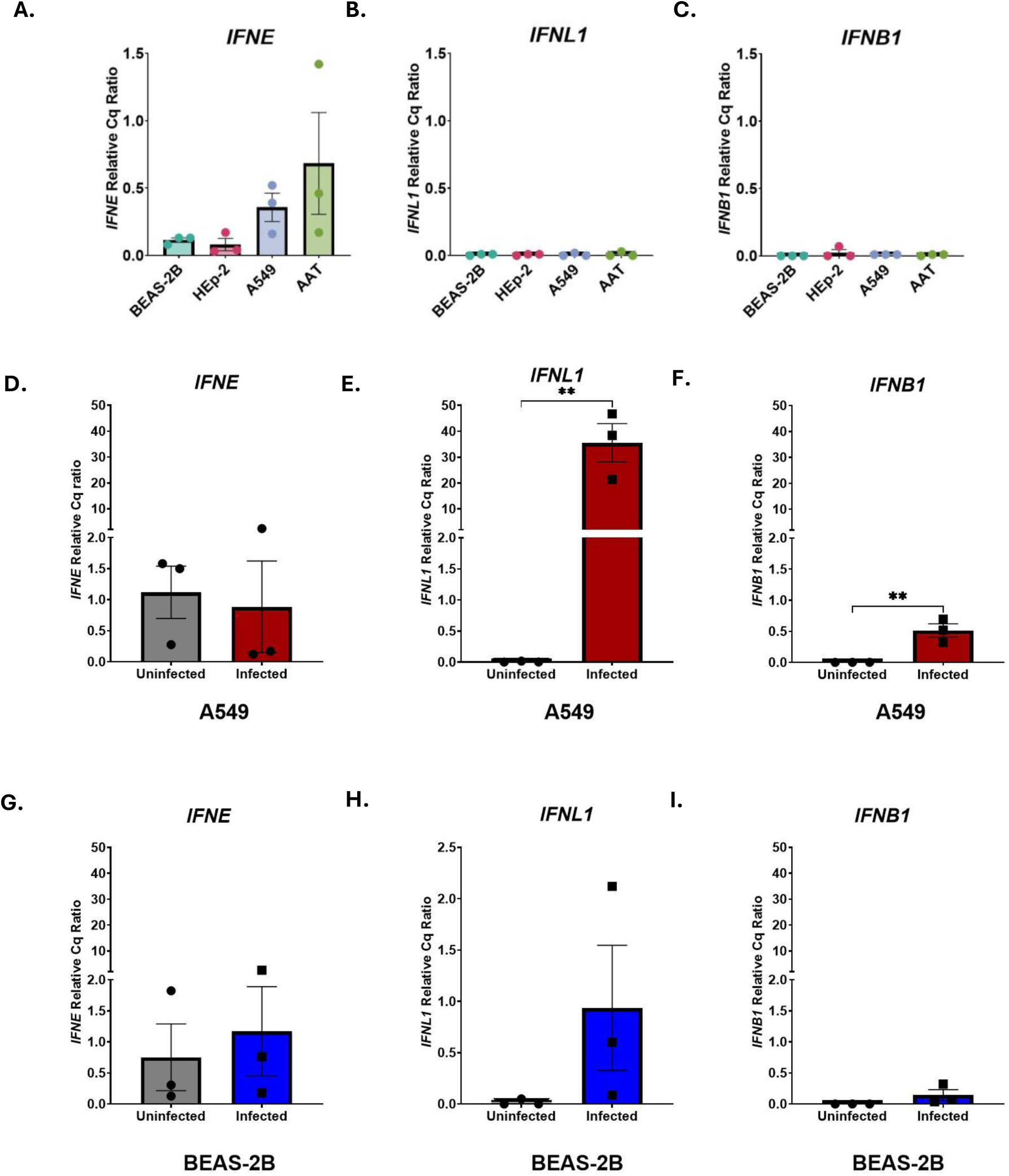
*IFNε* is endogenously expressed in immortalised airway epithelial cells. BEAS-2B, A549, AAT and HEp-2 cells were seeded in 24 well plate. 24 h post seeding total RNA was extracted, reverse transcribed and subjected to qPCR analysis from BEAS-2B, A549, AAT and HEp-2 cells for *IFNE*, *IFNL1* and *IFNB1* expression (ROCHE, Lightcycler 480). Graphs show relative ratio values for (**A**) *IFNE*, (**B**) *IFNL1* and (**C**) *IFNB1* normalised using *YWHAZ*, *IPO8* and *TBP* as house-keeping genes. Data are presented as mean of triplicate technical replicates (±SEM) for n=3 independent experiments. (**D-F**) A549 or (**G-I**) BEAS-2B cells were mock-infected or infected with RSV-A2/mKate2 (MOI 1). At 48 hpi total RNA was extracted, reverse transcribed and subjected to qPCR analysis of *IFNE*, *IFNL1* and *IFNB1* expression (ROCHE, Lightcycler 480). Graphs show relative ratio values for *IFNE*, *IFNL1* and *IFNB1* normalised using *YWHAZ*, *IPO8* and *TBP* as housekeeping genes. Data are presented as the mean of duplicate technical replicates (±SEM) for n=3 independent experiments at 48 hpi. Statistical analyses were conducted using an unpaired two-tailed Student’s t-test (* p<0.05, ** p<0.01).

### IFNε has antiviral activity against RSV that is concentration- and cell type-dependent

To assess if recombinant human IFNε (rhIFNε) protein can induce an antiviral effect within airway epithelial cell lines, A549 (**Figure 3A**) and BEAS-2B (**Figure 3B**) cells were pre-treated with rhIFNε, rhIFNβ1 and rhIFNλ1 protein (100 ng/mL) and infected with the highly IFN-sensitive virus encephalomyocarditis virus (EMCV) (MOI 1). IFNε pretreatment resulted in a 1.15-log_10_ reduction in EMCV titre versus the untreated in BEAS-2B cells (**Figure 3A**). This was similar to rhIFNλ1 pretreatment (1.7-log reduction) but much less effective than rhIFNβ1 pretreatment (3.5-log_10_ reduction). There was a non-statistically significant trend towards a reduction in virus titres within A549 cells (**Figure 3B**).

**Figure 3.**
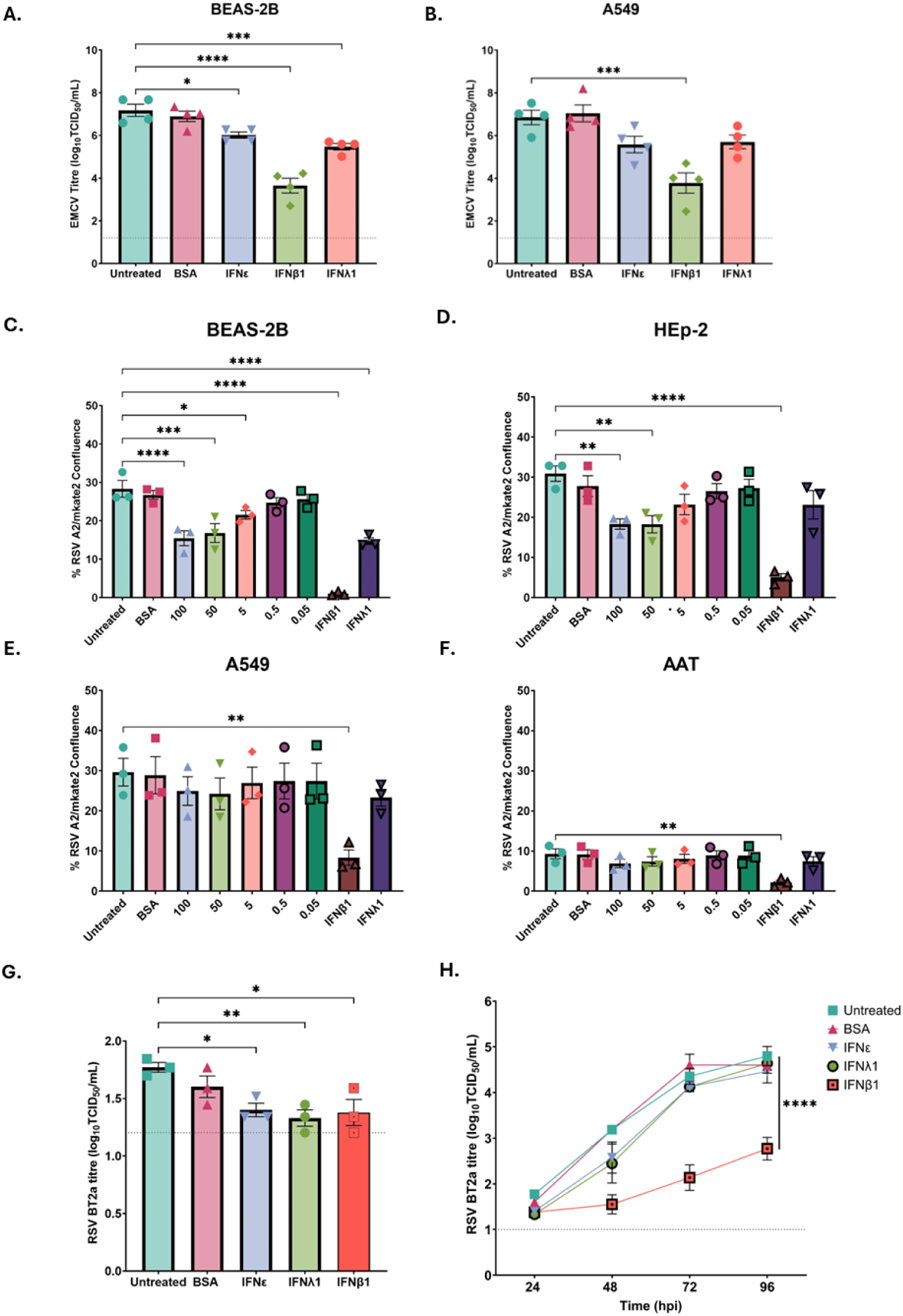
IFNε pretreatment reduces EMCV and RSV replication in BEAS-2B cells. **(A)** A549 and (**B**) BEAS-2B cells were pre-treated for 16 h with an IFN (100 ng/ml). Cells were then infected with EMCV (MOI=1) for 2 h at 37°C. Data are presented as mean of duplicate technical replicates (±SEM) for n=4 independent experiments. Statistical analysis was conducted using one-way ANOVA. (**C**) BEAS-2B, (**D**) HEp-2 (**E**) A549, and (**F**) AAT cells were pre-treated for 16 h with IFNε (100, 50, 5, 0.5, and 0.05 ng/mL), IFNβ1 or IFNλ1 (100 ng/mL). Cells were then infected with RSV-A2/mKate2 (MOI=0.3). (C-F) At 24 hpi mean % fluorescence ratio was measured using the brightfield and red fluorescence channels of the Celigo Imaging Cytometer (Nexcelom). Data presented as % fluorescence ratio for mean of triplicate technical replicates (±SEM) n=3 independent experiments. Statistical analysis (C-F) was conducted using one-way ANOVA. (**G-H**) BEAS-2B cells were pre-treated for 16 h with 100 ng/mL of IFN. Cells were then infected with RSV BT2a (MOI=0.3). Supernatants were collected at 24, 48, 72, and 96 hpi and TCID_50_ assays were conducted to determine RSV titres. Data are presented as the mean of duplicate technical replicates (±SEM) log_10_ TCID_50_/mL at (**G**) 24 hpi and a (**H**) 24-96 hpi time course for n=3 independent experiments. Statistical analysis for (**G**) was conducted using one-way ANOVA and for (**H**) using one-way ANOVA of AUC (* p<0.05, ** p<0.01, ***p<0.001, ****p<0.0001).

To explore the antiviral activity of rhIFNε against RSV, BEAS-2B (**Figure 3C**), HEp-2 **(Figure 3D**), A549 (**Figure 3E)** and AAT (**Figure 3F**) cells were pretreated with rhIFNε at a range of doses (100, 50, 5, 0.5 and 0.05 ng/mL) and infected with RSV-A2/mKate2 (MOI=0.3). A range was selected, as prior to this study, there was no evidence of IFNε’s antiviral effect against respiratory viruses, including, but not limited to, RSV. At 24 hpi, the mean % fluorescence ratio was measured using the brightfield and red fluorescence channels of the Celigo Imaging Cytometer (Nexcelom). BSA pretreatment (5 µg/mL) was a negative control for the recombinant protein stabilisation carrier, and rhIFNβ1 and rhIFNλ1 pretreatment (100 ng/mL) were used as positive controls to ensure cell responsiveness to IFNs. Using fluorescence as a surrogate marker of viral replication, rhIFNε pretreatment resulted in a significant reduction in virus fluorescence in BEAS-2B (**Figure 3C**) and HEp-2 cells (**Figure 3D**), but not in A549 or AAT cells (**Figure 3E-F**), although there was a trend towards reduction in the latter cell lines. Reduction in virus replication was dose-dependent, with 100 ng/mL of rhIFNε resulting in the greatest reduction compared to untreated controls (1.8-fold reduction) in the most responsive cell line (BEAS-2B) (**Figure 3C**). This was less potent than rhIFNβ1 (27-fold reduction) but similar to rhIFNλ1 (1.9-fold reduction) (**Figure 3C**). RSV-A2/mKate2 fluorescence was much lower in AAT cells (**Figure 3F**) compared to their parental counterpart A549 cells. To assess the antiviral effect against RSV BT2a [33], BEAS-2B cells were pre-treated with 100 ng/mL of rhIFNε, rhIFNβ1and rhIFNλ1 and infected with RSV BT2a (MOI 0.3). BEAS-2B cells were selected as they are a non-tumour transformed cell line and more responsive to IFN stimulation. We found a 0.37-Log_10_ reduction at 24 hpi (**Figure 3G**) in RSV titres in rhIFNε pre-treated cells compared to untreated controls. A similar reduction was seen with rhIFNβ1 (0.39-Log_10_ reduction) and rhIFNλ1 (0.44-Log_10_ reduction) pre-treatment at 24 hpi. Thereafter, while mean virus growth kinetics were delayed slightly at 48 hpi, titres at 72 and 96 hpi were indistinguishable from untreated- or BSA-controls (**Figure 3H**).

### Relative antiviral activity of IFNε, IFNλ1 and IFNβ1 against RSV

To further facilitate meaningful comparisons of the relative antiviral activity of rhIFNε with rhIFNβ1 and rhIFNλ1, we determined the half-maximal inhibitory concentration (IC_50_) of these IFNs against RSV in BEAS-2B cells. Cells were pre-treated with decreasing concentrations of rhIFNε (**Figure 4A**), rhIFNλ1 (**Figure 4B**) and rhIFNβ1 (**Figure 4C**) and infected with RSV-A2/mKate2 (MOI=0.1). As anticipated, IFNε possessed the highest best-fit IC_50_ value (9 ng/mL), followed by IFNλ1 (2.8 ng/ml) and IFNβ1 (0.24 ng/mL).

**Figure 4.**
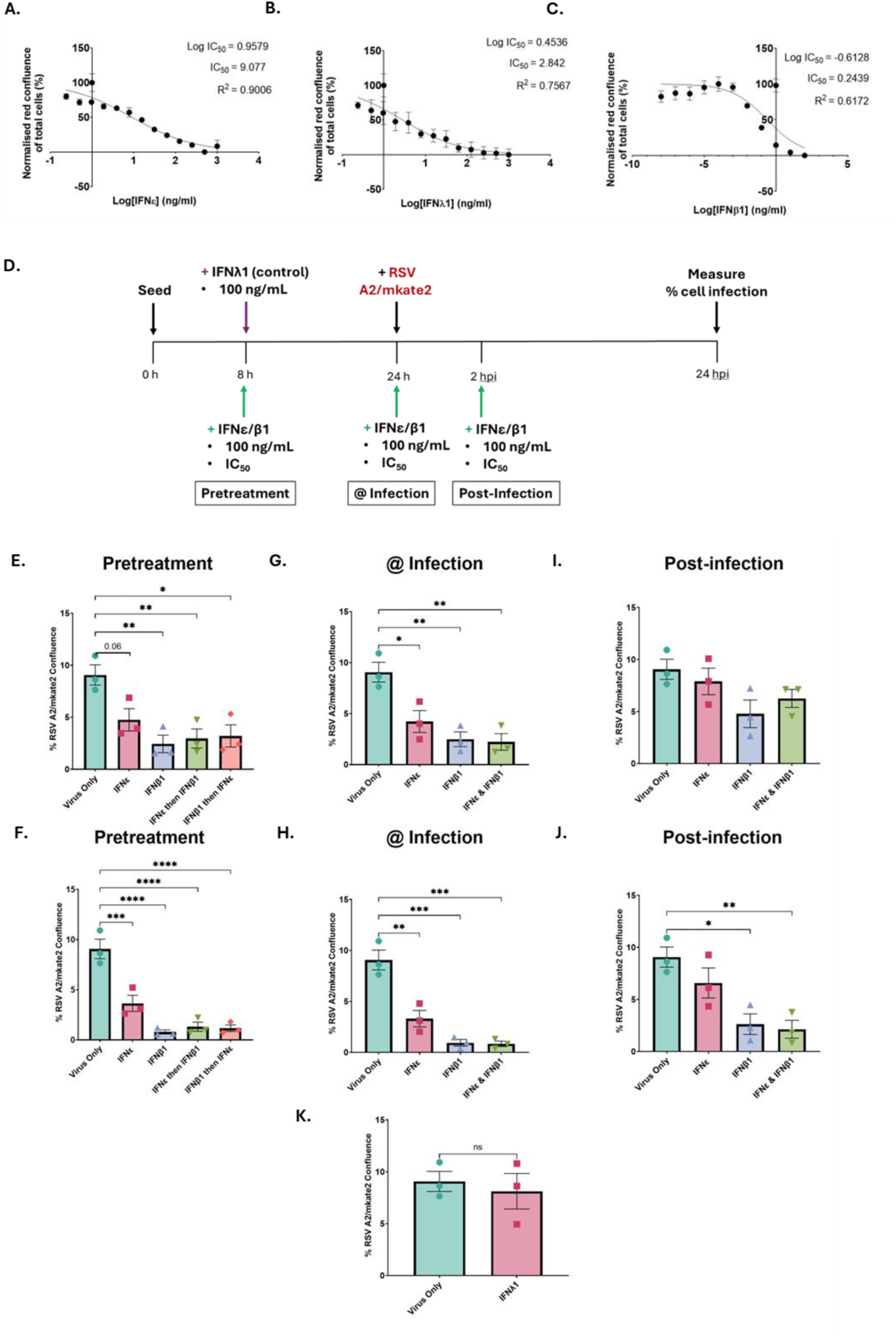
IFNε possesses an increased IC_50_ concentration against RSV compared to IFNβ1. BEAS-2B cells were pre-treated for 16 h with decreasing concentrations of (**A**) IFNε, (**B**) IFNλ1 and (**C**) IFNβ1. Cells were then infected with RSV-A2/mKate2 (MOI=0.1). At 24 hpi mean % fluorescence ratio was measured using the brightfield and red fluorescence channels of the Celigo Imaging Cytometer (Nexcelom). Statistical analysis (**A-C**) was conducted fitting a non-linear dose-response curve for mean of triplicate technical replicates for n=2 (IFNε and IFNλ1) or n=3 (IFNβ1) independent experiments in triplicate. (**D**) Summary of Experimental set-up for pretreatment, treatment at infection and treatment post-infection experiment. *IFNLR1* Cas9 knockout BEAS-2B cells were either pretreated (**E-F**), treated at infection (**G-H**) or treated 2 hours post infection (**I-J**) with the IC_50_ concentration (**E**, **G**, **I**) or 100 ng/mL (**F**, **H**, **J**) of recombinant human IFNε and IFNβ1 protein. Cells were infected with RSV-A2/mKate2 (MOI=0.1). (**K**) *IFNLR1-/-* BEAS-2B cells pretreated with 100 ng/mL of recombinant human IFNLR1. Data presented as % fluorescence ratio for as the mean of triplicate technical replicates (±SEM) n=3 independent experiments. Statistical analysis was conducted using one-way ANOVA (* p<0.05, **p<0.01, ***p<0.001, ****p<0.0001).

Given the limited understanding of how IFNε exerts its antiviral activity, we aimed to confirm that its presence would not interfere with the activity of other type I IFNs, specifically IFNβ1, in our study. Therefore, to further explore this without the interference from type III IFNs, *IFNLR1*^-/-^ BEAS-2B cells (**Figure 4D**) were either pre-treated (**Figure 4E-F**), treated at infection (**Figure 4G-H)** or at 2 h post-infection (**Figure 4I-J**) with the IC_50_ concentrations (**Figure 4E, G, I**) or 100 ng/mL (**Figure 4F, H, J**) of rhIFNε and rhIFNβ1. Cells were infected with RSV-A2/mKate2 (MOI=0.1). At 24 hpi, IC_50_ concentrations of rhIFNε and rhIFNβ1 resulted in a reduction in viral spread with pre-treatment and with treatment at infection (**Figure 4E-H**). However, IC_50_ concentrations of rhIFNε and rhIFNβ1 were not sufficient to significantly diminish RSV spread when treatment commenced 2 hpi (**Figure 4I)**. When pre-treated with both IFNs sequentially (rhIFNε and then rhIFNβ1 or vice versa) or treated with a combination of the two IFNs at the same time, the antiviral activity of rhIFNβ1 was not altered in the presence of rhIFNε. The greatest reduction in virus spread was seen when rhIFNβ1 was present regardless of the sequence of addition. Cells pre-treated or treated at infection with 100 ng/mL resulted in robust inhibition of viral spread, with those treated with 100 ng/mL rhIFNβ1 showing the greatest reduction (**Figure F-H**). Only 100 ng/mL of rhIFNβ1 inhibited viral spread significantly with treatment post-infection (**Figure 4J**). rhIFNλ1 pre-treatment (100 ng/mL) in *IFNLR1^-/-^* BEAS-2B cells did not result in a significant reduction in virus spread (**Figure 4J**). These results demonstrated that rhIFNε does not interfere with the antiviral activity of rhIFNβ1, despite both binding to the same receptor, at IC_50_ concentrations or 100 ng/mL.

### IFNε induces antiviral activity against Sendai Virus but not SARS-CoV-2

To assess if an antiviral effect was evident beyond EMCV and RSV, A549 (**Figure 5A**) and BEAS-2B (**Figure 5B**) cells were pre-treated with a dose range of rhIFNε (100, 50, 5, 0.5 and 0.05 ng/mL) and infected with rSeV/eGFP (MOI=0.3). No significant reduction was observed in A549 cells, despite a reduction trend being observed (**Figure 5A**). At 24 hpi, a similar dose-dependent antiviral activity of rhIFNε was evident against rSeV/eGFP, as previously described for RSV A2/mkate2 in BEAS-2B cells (**Figure 5B**). The antiviral effect of rhIFNε pre-treatment (2.2-fold reduction) was like that of rhIFNλ1 (4-fold reduction) in BEAS-2B cells but less impressive compared to rhIFNβ1 (16.3-fold reduction) (**Figure 5B**). In contrast, AAT cells pretreated with rhIFNε, rhIFNβ1 or rhIFNλ1 (100 ng/mL) and infected with the SARS-CoV-2 Delta variant (MOI=0.3), had no significant reductions in viral replication with IFNε pre-treatment, a slight but significant reduction in virus replication with rhIFNλ1 pre-treatment, and a complete abrogation of virus replication following rhIFNβ1 pre-treatment at 48 hpi (**Figure 5C**) and over a time course of infection (**Figure 5D**). These results suggest that rhIFNε possesses antiviral activity against some RNA viruses, but not SARS-CoV-2 Delta Variant, at least at the dose tested, and its potency is clearly reduced compared to rhIFNβ1.

**Figure 5.**
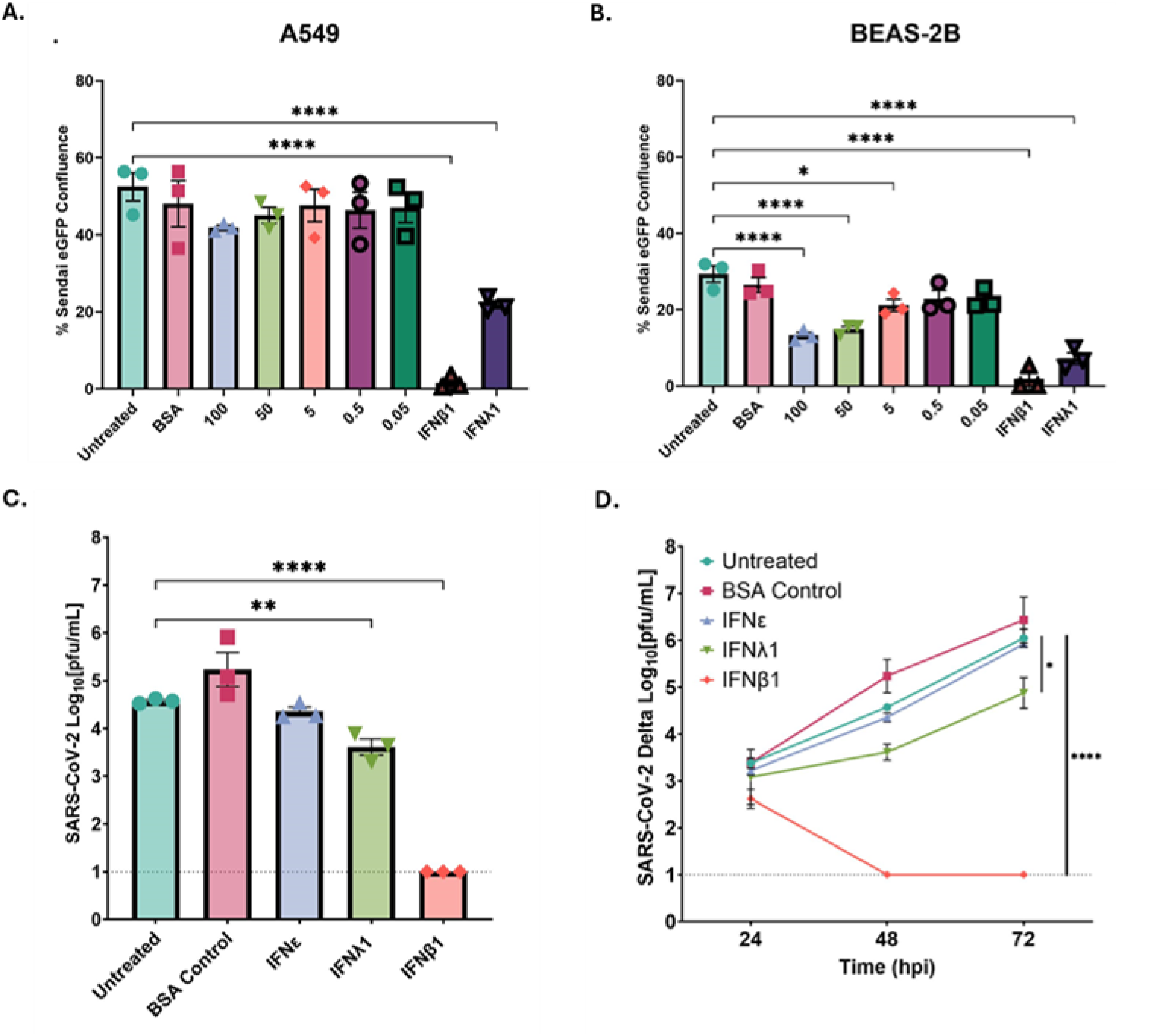
Relative potency of IFNε, IFNλ1 and IFNβ1 pre-treatment against Sendai virus and SARS-CoV-2. (**A**) A549 and (**B**) BEAS-2B cells were pre-treated for 16 h with IFNε (100, 50, 5, 0.5, and 0.05 ng/ml), rhIFNβ1 or rhIFNλ1 (100 ng/mL). Cells were infected with rSeV/eGFP (MOI=0.3). At 24 hpi mean % fluorescence ratio was measured using the brightfield and green fluorescence channels of the Celigo Imaging Cytometer (Nexcelom). Data presented as % fluorescence ratio for mean of triplicate technical replicates (±SEM) for n=3 independent experiments. Statistical analysis was conducted using one-way ANOVA. (**C-D**) AAT cells were pre-treated for 16 h with rhIFNε, rhIFNλ1 and rhIFNβ1 (100 ng/mL). Cells were then infected with SARS-CoV-2 Delta variant (MOI=0.3). Supernatants were collected at 24, 48, and 72 hpi and plaque assays were performed on VATs. Data presented as the mean of triplicate technical replicates (±SEM) log_10_ pfu/mL at (**C**) 48 hpi and a (**D**) 24-72 hpi time course for n=3 independent experiments. Statistical analysis (**C**) was conducted using one-way ANOVA (**D**) Statistical analysis was conducted using one-way ANOVA of AUC (* p<0.05, **p<0.01, ***p<0.001, ****p<0.0001).

### JAK-STAT inhibitor alters IFNε-mediated reduction of viral replication

To confirm that IFNε was signalling via the classical type I IFN signal induction pathway (IFNAR-JAK-STAT) in airway epithelial cells, BEAS-2B (**Figure 6A-B**) and A549 (**Figure 6C-D**) cells were pre-treated with the JAK inhibitor Ruxolitinib for 2 h prior to rhIFNε (**Figure 6A, C**) and rhIFNβ1 (**Figure 6B, D**) pre-treatment (100 ng/mL). Pre-treatment with Ruxolitinib eliminated the reduction in viral replication with rhIFNε pretreatment in BEAS-2B cells (**Figure 6A**). A similar trend was seen for rhIFNβ1-pre-treated BEAS-2Bs (**Figure 6B**), although the concentration of Ruxolitinib used was not sufficient to completely block rhIFNβ1 signalling at 100 ng/mL. A549 cells did not produce a significant reduction in viral replication with rhIFNε treatment as described above (**Figure 6C)**. However, pre-treatment of A549 cells with 100 ng/mL of rhIFNβ1 or rhIFNβ1 did result in a significant reduction in viral replication (**Figure 6D**). These results confirm the increased potency of rhIFNβ1 compared to rhIFNε at 100 ng/mL but that, like rhIFNβ1, rhIFNε utilises the JAK-STAT pathway to exert its antiviral effect against RSV.

**Figure 6.**
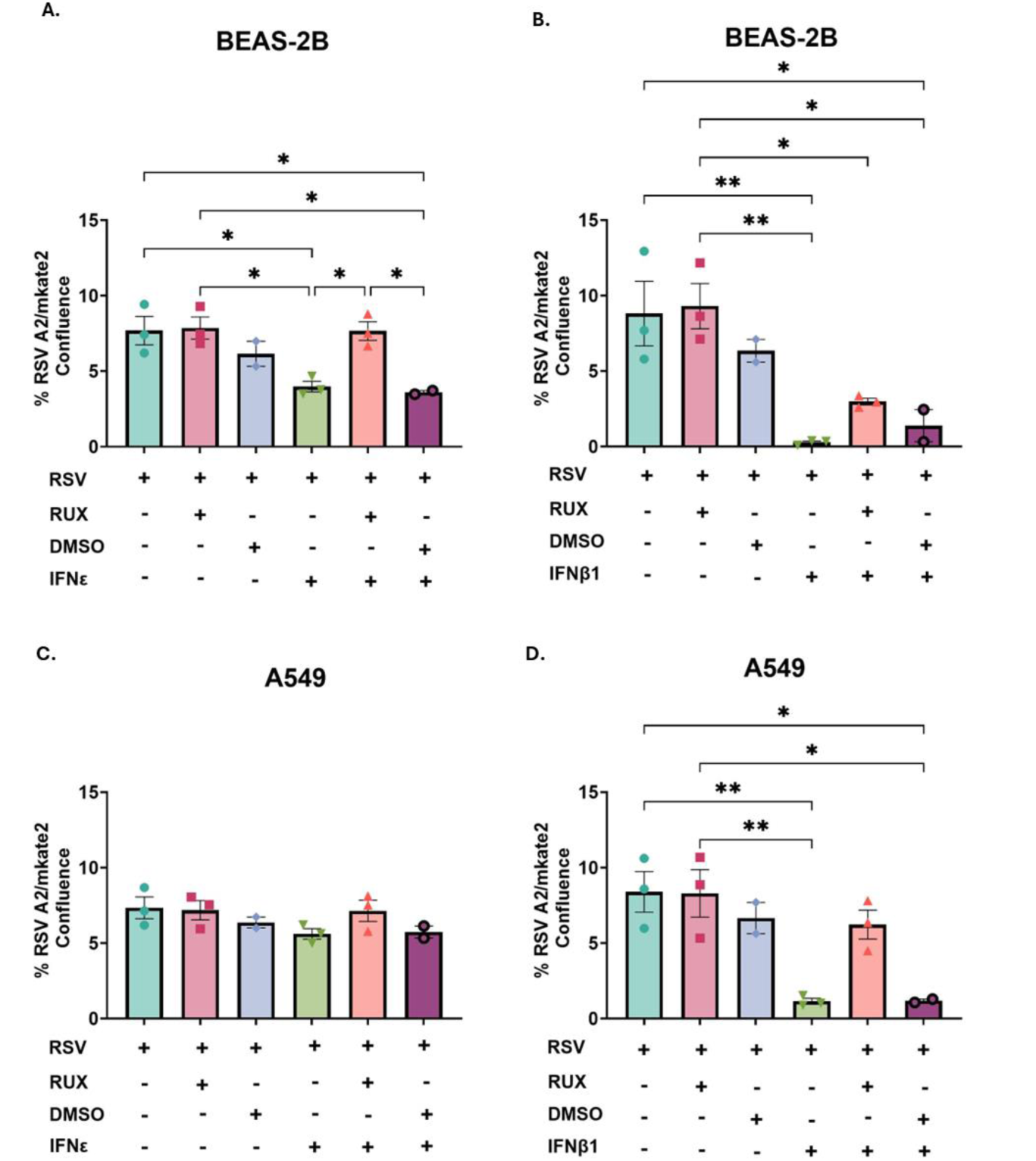
JAK-STAT inhibitor alters IFNε-mediated reduction of viral replication. BEAS-2B and A549 cells were pre-treated with Ruxolitinib (INCB018424, Selleck Chemicals) or DMSO (2.5 μM) for 2 h before 16 h IFNε (**A-B**), and IFNβ1 (**C-D**) pre-treatment (100 ng/mL). Cells were then infected with RSV (RSV-A2/mKate2) (MOI=0.1). At 24 hpi BEAS-2B (**A**, **C**) and A549 (**B**, **D**) cells mean % fluorescence ratio was measured using the brightfield and red fluorescence channels of the Celigo Imaging Cytometer (Nexcelom). Data presented as % fluorescence ratio for mean of triplicate technical replicates (±SEM) for n=3 independent experiments for RSV, RSV+RUX, RSV+IFNε and RSV+RUX+IFNε. Data presented as % fluorescence ratio of triplicate technical replicates for (±SEM) n=2 independent experiments for RSV+DMSO and RSV+DMSO+IFNε. Statistical analysis was conducted using one-way ANOVA (* p<0.05 **p<0.01).

### Differential ISG expression kinetics following rhIFNε, rhIFNλ1, and rhIFNβ1 treatment

In our hands, IFNε was capable of inhibiting RSV replication only at early time points post-infection suggesting that the expression of downstream antiviral restriction factors may be transient. To assess if any differences were evident in the magnitude and expression kinetics of downstream antiviral restriction factors with rhIFNε treatment compared to rhIFNβ1 and rhIFNλ1, RT-qPCR was conducted on RNA extracted from BEAS-2B cells treated with rhIFNε, rhIFNλ1 and rhIFNβ1 (100 ng/mL) (**Figure 7A-F**). rhIFNε induced well known antiviral proteins, such as *MX1* (**Figure 7A**), *ISG15* (**Figure 7B**), *IFIT1* (**Figure 7C**) and *RSAD2* (**Figure 7D**). Similarly, IFNε also induced type I IFN-associated pro-inflammatory genes, such as *IRF1* (**Figure 7E**), and *CXCL10* (**Figure 7F**), which were previously reported to be induced only at low levels and transiently by rhIFNλ1 or other type III IFNs [38]. rhIFNε possessed unique ISG expression kinetics, with expression peaking early (4 h post-treatment), in comparison to rhIFNβ1, which induced a more potent and enduring response, and rhIFNλ1, which induced a slower yet incrementally increasing response through 24 h post-treatment. These results demonstrate that rhIFNε can induce a typical pro-inflammatory type I IFN antiviral gene signature in airway epithelial cells but that temporal expression of downstream ISGs is transient compared to rhIFNβ1 and rhIFNλ1 treatment.

**Figure 7.**
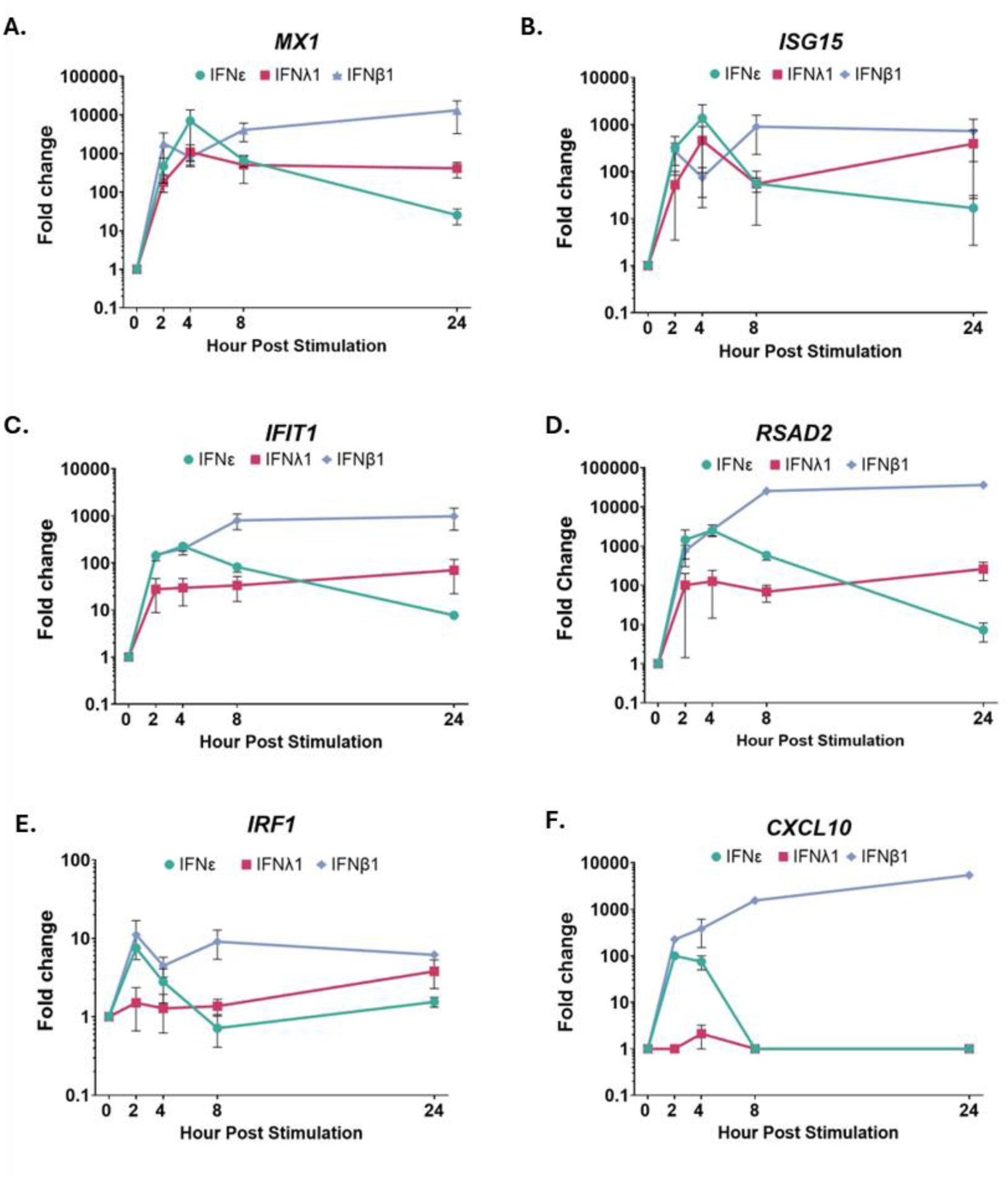
Differential induction of antiviral and pro-inflammatory ISGs by IFNε. BEAS-2B cells were treated with IFNε, IFNλ1 or IFNβ1 (100 ng/ml) and total RNA was extracted immediately from untreated (t=0) and at indicated time points post-stimulation. RNA was subjected to RT-qPCR analysis of *IFIT1*, *ISG15*, *MX1, RSAD2, IRF1 and CXCL10* expression (ROCHE, Lightcycler 480). Graphs show relative ratio values for antiviral ISGs (**A**) *MX1*, (**B**) *ISG15*, (**C**) *IFIT1,* (**D**) *RSAD2 and pro-inflammatory ISGs* (**E**) *IRF1* and (**F**) CXCL10, normalised using *YWHAZ*, *IPO8* and *TBP* as housekeeping genes. Data presented (**A**-**D**) are mean of duplicate technical replicates (±SEM) fold change from mock for n=3 independent experiments and data presented (**E**-**F**) are mean of duplicate technical replicates (±SEM) fold change from mock for n=2 independent experiments.

## Discussion

Conflicting data exists surrounding the role of IFN responses during RSV infection, particularly among those most vulnerable to severe RSV-related disease, infants aged 6 weeks to 6 months of life. Most of what is known describe the “classical” type I (IFNβ1 and IFNα’s) and type III (IFNλ1-4) IFNs [39, 40]. However, little is understood about the “non-classical” IFN subtypes, such as IFNε. Previously we exploited a unique cohort of primary paediatric airway epithelial cell samples derived from the same healthy infants at birth and 1 year, to explore the development of innate immune responses to RSV infection in the first year of life (unpublished data, Groves HE, McCabe M, Coey J, Broadbent L, Guo-Parke H, Lopez Campos G, et al). We present here the differential expression of *IFNE* with chronological age, between healthy newborn and 1-year infants, both endogenously and following RSV infection in WD-PNECs cultures. This timespan coincides with those infants with high susceptibility rates to severe RSV-related disease (newborn) and those with lower frequencies of severe infection (1 year). Previous studies have reported endogenous and infection-induced *IFNE* expression in both the upper and lower respiratory tracts in humans and other mammals, with variations in expression linked to age and disease severity [24, 28, 29, 41, 42]. IFNε is conserved across various mammalian species [24, 43–46] but is a pseudogene in species susceptible to infections that are relevant to this study [47–49], alluding to a evolutionarily significant mucosal function. Collectively, these data suggest that IFNε may play a protective role in the respiratory epithelium during early-life viral infection and elevated levels in infants may be an indication of an improved ability to curtail and/or resolve infection.

As epithelial cells are the primary target of RSV infection *in vivo*, we assessed the expression of *IFNE* in immortalised epithelial cell lines commonly used to model RSV infection. *IFNE* was endogenously expressed at low levels in all cell lines tested. Recently, Martínez-Espinoza *et al* described increased *IFNE* expression during infection with RSV and hMPV, with peak *IFNE* expression at 48 hpi in A549 cells for both viruses [41]. We did not see this increase in *IFNE* expression with RSV infection. This discrepancy may be due to various factors, including differences in cell culturing conditions, viral strains used, assays performed, and other experimental variables between authors. Our observations are consistent with its reported unstimulated endogenous expression within the epithelium of the FRT and male reproductive tract in humans and mice [21, 22, 50–52]. However, we did observe an increase in *IFNE* expression within our RSV-infected WD-PNEC cultures, mirroring the increase Martinez-Espinoza *et al* reported within human bronchial airway epithelial cells 3-5 days post RSV-infection [41]. Notably, Martínez-Espinoza *et al* suggested that RSV and hMPV induced *IFNE* expression in A549 cells via RIG-I, with contributions from MDA5 [41]. This suggests that viral RNA plays a role in the induction of *IFNE* during active infection. The endogenous expression of *IFNE* is conserved across multiple mucosal sites, concentrated primarily in the epithelial cells lining these surfaces. Consequently, it may be regulated by distinct stimuli in a cell/tissue-specific manner, possibly by distinct cell populations in these sites which fluctuate with tissue-specific stimuli. However, no studies have been published regarding IFNε’s role in human respiratory epithelium on a single-cell level, and further work needs to be conducted regarding *IFNE* induction and regulation in more physiologically relevant primary human airway epithelial models.

As a result of *IFNE’s* endogenous epithelial expression, further investigation was prompted regarding its possible contribution to localised antiviral activity. We showed that rhIFNε provided antiviral protection against EMCV infection in BEAS-2B and A549 cells, consistent with similar antiviral effects observed in cervical WISH cells [25]. Martínez-Espinoza *et al* demonstrated the antiviral activity of rhIFNε against RSV and hMPV in A549 cells [41], which were shown to be highly permissive to RSV infection [53]. Therefore, we chose to assess the antiviral properties of rhIFNε in multiple respiratory cell lines. Pre-treatment with rhIFNε resulted in a significant dose-dependent reduction in BEAS-2B and HEp-2 cell lines. BEAS-2B cells were observed to express high levels of transcription factors and antiviral ISGs upon RSV infection and multiple IFN subtype treatment, indicating a strong response to rhIFNε [53]. We did not observe a significant reduction in virus fluorescence in A549 and AAT cells, when treated with 100 ng/mL of rhIFNε, consistent with Martínez-Espinoza *et al*., who found that at least 250 ng/mL was needed for a significant effect on RSV fluorescence or titres in A549 cells [41]. In addition, prototypic lab-adapted strains, such as RSV A2/mkate2, have been reported to result in greater cytopathogenesis and induce greater concentrations of proinflammatory cytokines in the respiratory epithelium [33]. Therefore, we also chose to assess the antiviral activity of rhIFNε against a low passage clinical isolate of RSV, which more accurately reflects RSV infection *in vivo*. We found that IFNε pretreatment of BEAS-2B cells was antiviral against the clinical isolate RSV BT2a, with a significant reduction in infectious viral release at 24 hpi. However, without being replenished, this antiviral effect was quickly lost as the infection progressed. BEAS-2B cells were selected as the representative cell line due to their significant reduction in RSV A2/mKate2 infection following 100 ng/mL rhIFNε pretreatment and their non-cancerous immortalized state. Our data emphasizes the importance of cell line selection in studying viral IFN sensitivity, especially for lesser-understood IFN subtypes and suggests that while IFNε may not be the primary defender against viral infection in epithelial cells, they support its potential role as an endogenous antiviral mucosal defence cytokine.

The relative antiviral activities of each IFN were reflected in their IC_50_ values. Our data indicated that rhIFNε was ∼3-fold less potent than rhIFNλ1 and ∼38-fold less potent than rhIFNβ1, similar to those described for recombinant mouse IFNε (rmIFNε) against EMCV in WISH cells [25]. Interestingly, the IC_50_ concentration of rhIFNε and 100 ng/mL was sufficient to reduce viral replication under pretreatment and treatment-at-infection conditions, but not with treatment 2 h post RSV infection in BEAS-2B cells. However, its inability to provide protection against infection when added post-RSV infection most likely reflects its reduced antiviral potency and RSV’s recognised ability to efficiently modulate IFN responses [54]. This is consistent with data showing a reduction in Zika virus replication in HRT8 cells with rmIFNε pretreatment, but not post-infection [50]. As expected, RSV was also able to antagonise IFNβ1’s antiviral activity when added post-infection at its IC_50_ concentration, as IFNε and IFNβ1 share the same signalling pathway. However, when added 2 h post-RSV infection at 100 ng/mL rhIFNβ1 was able to reduce viral replication significantly, most likely reflecting rhIFNβ1’s greater potency. As anticipated the antiviral activity of rhIFNβ1 consistently superseded rhIFNε when added before, after or simultaneously with rhIFNε. Like IFNβ1, IFNε possesses a greater affinity for the IFNAR1 chain [25]. However, IFNε binds to IFNAR1 at a much lower affinity than IFNβ1 (100-1000-fold), which may partly explain the increased antiviral capacity of IFNβ1 in this context [25].

Infection with the related virus rSeV/eGFP confirmed that IFNε’s antiviral effect was not restricted to RSV, with the greatest reduction in rSeV/eGFP fluorescence also observed within BEAS-2B cells. However, we did not observe an antiviral effect against the SARS-CoV-2 Delta variant in AAT cells at the very high dose tested (100 ng/mL), a variant that was previously described to have increased sensitivity to IFNs [55–58]. To our knowledge, this is the first report demonstrating SARS-CoV-2’s insensitivity to rhIFNε [59]. Although our results suggest that rhIFNε was not effective in altering SARS-CoV-2 infection, it is important to note that we used a modified immortalized cell line highly susceptible to SARS-CoV-2 infection. Further studies in a more physiologically relevant nasal epithelial model are required, given evidence that IFNε may influence age-related responses to SARS-CoV-2 [29, 30]. It is possible that during SARS-CoV-2 infection, IFNε possess functions beyond direct antiviral activity, as suggested by its wider immunomodulatory functions [60]. Overall, our data demonstrate that respiratory viruses possess differential sensitivities to rhIFNε restriction and emphasise the importance of assessing the antiviral capacity against multiple virus families.

Our data using a JAK-STAT signalling inhibitor confirmed that IFNε signals through this pathway to exert its antiviral activity against RSV. These data provided the rationale to explore the magnitude and kinetics of the induction of downstream ISGs with rhIFNε treatment. Recently, IFNε was shown to induce common ISGs upregulated during RSV infection [41]. Our data showed that rhIFNε treatment induced a unique ISG expression pattern with expression of four antiviral ISGs associated with RSV infection (*MX1*, *ISG15*, *IFIT1* and *RSAD2*) peaking at 4 h post-stimulation. IFNε’s transient induction pattern was consistent with that observed against ZIKV infection in human FRT cell lines [50] and this rapid transient ISG induction may be reflective of its short-term reduction in viral titre at 24 hpi in BEAS-2B cells, but not at later time points. Furthermore, rhIFNε, like rhIFNβ1, induced the expression of the transcription factor *IRF1* and the chemokine *CXCL10*. rhIFNλ1 treatment did not induce *IRF1* or *CXCL10* expression to the same magnitude compared to rhIFNε and rhIFNβ1. Rapid increased *IRF1* and *CXCL10 expression* are considered type I IFN-associated [38]. However, type III IFNs were shown to induce *IRF1* and *CXCL10* expression at lower levels [38]. Together our results demonstrated that rhIFNε can induce a strong classical type I antiviral gene signature in airway epithelial cells but its potency and expression kinetics are distinct compared to rhIFNβ1 and rhIFNλ1.

IFNε’s unique low-level endogenous expression, its reduced potency, and transient ISG expression kinetics, may strike a balance between the beneficial effects of type I IFN and minimizing the tissue toxicity associated with its presence [61]. The question remains whether the endogenous expression of IFNε results in a continuously primed airway epithelium, creating an endogenous barrier to infection. If so, IFNε or the ISGs it induces could be leveraged as potential biomarkers to identify those children at increased risk of severe disease in which no high-risk factors have been identified (6 weeks and 6 months). Could the unique properties of IFNε be further harnessed as an antiviral to non-destructively boost protective responses to infection? Further work is required to explore IFNε’s ability to boost or improve innate immune responses within primary airway epithelial cell cultures.

## Conclusion

IFNε is a highly conserved type I IFN expressed at multiple mucosal surfaces and is recognised for its antimicrobial abilities in the FRT. We demonstrated differential expression of *IFNE* with age and RSV infection in human nasal epithelial cells and confirmed its antiviral activity against a clinical isolate of RSV. Although less potent than rhIFNβ1 or rhIFNλ1, rhIFNε’s unique ISG expression kinetics at 100 ng/mL and mild inflammatory profile suggest it may play a role in mitigating RSV infection in airway epithelium. Reduced IFNε expression in newborns may partially explain their heightened vulnerability to RSV, presenting an opportunity for therapeutic interventions. Further studies are needed to explore IFNε’s potential as an antiviral therapy or prognostic tool for early-life respiratory infections.

## CRediT authorship contribution statement

**Mary McCabe**: Writing -Original Draft, Validation, Methodology, Investigation, Formal Analysis, Data Curation. **Helen E. Groves**: Conceptualization, Methodology, Supervision. **Erin Getty**: Investigation, Writing – Review & Editing. **Emma Campbell**: Investigation. **Connor G.G Bamford**: Methodology, Resources, Writing – Review & Editing, Supervision. **Guillermo Lopez Campos**: Methodology, Conceptualization, Writing – Review & Editing, Supervision. **Michael D. Shields**: Conceptualization, Writing – Review & Editing, Funding acquisition. **Ultan F. Power**: Conceptualisation, Resources, Writing – Review & Editing, Supervision, Funding acquisition.

## Acknowledgements

This work was supported by Wellcome Trust grant number 104516/Z/14/Z to HG; PHA HSCNI R&D Division funding (grant numbers RRG/3244105 and Com/4044/09), and Department for Economy, Northern Ireland to UFP.

